# Dynamic changes in natural killer cell subset frequencies in the absence of cytomegalovirus infection

**DOI:** 10.1101/745190

**Authors:** Ivayla E. Gyurova, Heinrich Schlums, Heidi Sucharew, Lilliam Ambroggio, David E. Ochayon, Hannah Than Win, Yenan T. Bryceson, David I. Bernstein, Stephen N. Waggoner

## Abstract

Individuals lacking functional natural killer (NK) cells suffer severe, recurrent infections with cytomegalovirus (CMV), highlighting the critical role of NK cells in antiviral defense. Therefore, ongoing attempts to develop an efficacious vaccine to prevent CMV infection should potentially aim to elicit NK-cell antiviral responses as an accessory to conventional T- and B-cell based approaches. In this regard, CMV infection provokes marked phenotypic and functional differentiation of the NK-cell compartment, including development of adaptive NK cells that exhibit enhanced antiviral activity. We examined longitudinal blood samples collected from 40 CMV-seronegative adolescents to ascertain whether a CMV glycoprotein B (gB) vaccine in the absence of CMV infection can stimulate differentiation or expansion of CMV-associated subsets of NK cells. Study participants uniformly lacked the CMV-dependent NKG2C^+^ subset of NK cells, suggesting that an adjuvanted CMV gB vaccine alone is an inadequate stimulus for sustained expansion of these cells. In contrast, we observed unexpected dynamic fluctuations in the frequency of NK cells lacking FcRγ, EAT-2, and SYK, which were independent of vaccination or CMV infection. Whereas FcRγ ^neg^ NK cells in CMV infection are reported to express increased levels of the maturation marker CD57, the FcRγ^neg^ NK cells observed in our CMV-negative vaccine cohort express less CD57 than their FcRγ^+^ counterparts. The FcRγ^neg^ NK cells in CMV-negative individuals were also functionally distinct from this subset in CMV infection, exhibiting comparable IFN-γ production and degranulation as FcRγ^+^ NK cells in response to cytokine or antibody-dependent stimuli. These results suggest that frequencies of some NK cell subsets may increase in response to unknown environmental or inflammatory cues distinct from that which occurs after CMV infection. Greater understanding of the nature of the signals driving CMV-independent accumulation of these subsets should permit development of mechanisms to facilitate vaccine-driven expansion of CMV-reactive NK cells.

## Introduction

Cytomegalovirus (CMV) is a significant global cause of morbidity with manifestations of infection ranging from subclinical disease to death. Congenital infection and the infection of immunocompromised patients, including transplant recipients, result in the most severe consequences of CMV in the human population. Congenital CMV accounts for roughly 400 deaths and more than 5,000 developmentally impaired children each year in the United States (Dollard et al., 2007). Therefore, effective strategies to prevent or control infection are desperately needed.

Unfortunately, CMV has proven to be a challenging target for vaccine development. To date, most CMV vaccine efforts focus on elicitation of antibodies against viral glycoproteins or generation of antiviral T-cell responses (Schleiss et al., 2017). Administration of a MF59-adjuvanted CMV glycoprotein B (gB) vaccine to CMV-seronegative adolescent girls induced strong gB-specific antibody responses and afforded 43% protective efficacy (Bernstein et al., 2016). The same vaccine conferred short-lived, 50% protection, against CMV infection in seronegative post-partum women (Pass et al., 2009) and reduced post-transplant viral load when given to patients awaiting a kidney or liver transplant (Griffiths et al., 2011). While promising, these results indicate that humoral responses against gB may be insufficient to effectively prevent CMV infection in many individuals. DNA vaccines aimed at eliciting CMV-reactive T cells have also afforded minimal protection in a transplant patient-based clinical trial (Vincenti et al., 2018). These advances prompted development of new vaccines aimed at eliciting both humoral and cellular immunity (John et al., 2018), but it remains unclear whether other arms of the immune response must be engaged to effectively prevent CMV infection.

Natural killer (NK) cells are critical antiviral effectors that produce IFN-γ (Orange et al., 1995), lyse virus-infected cells (Santoli et al., 1978), and regulate adaptive immune responses (Crome et al., 2013; Welsh and Waggoner, 2013; Cook et al., 2014; Crouse et al., 2015; Pallmer and Oxenius, 2016; Schuster et al., 2016). NK cells play an important role in control of CMV infection in both mice and humans (Bukowski et al., 1983; Biron et al., 1989). Since NK cells lack the somatically rearranged antigen-specific receptors characteristic of T and B cells, and because they were previously thought to be short-lived cells (Zhang et al., 2007), vaccine triggering of NK cells has historically been considered of little value. However, recent data suggests that the innate immune system makes important contributions to vaccine-elicited protection against infection (Cerwenka and Lanier, 2016; Netea et al., 2016). Specifically, long-lived populations of adaptive NK cells with antigen-specific features similar to those of memory T cells have emerged as potential new targets of vaccines aimed at preventing CMV infection (Sun and Lanier, 2009; Cooper and Yokoyama, 2010; Paust and von Andrian, 2011; Vivier et al., 2011).

Immunological memory in virus-specific NK cells is widely described in the context of murine CMV. In C57BL/6 mice, a mouse CMV gene product engages an activating NK cell receptor, Ly49H (*Klra8*), promoting clonal expansion and contraction of the Ly49H-expressing subset of NK cells (Brown et al., 2001; Daniels et al., 2001; Dokun et al., 2001; Lee et al., 2001; Arase et al., 2002). Thereafter, a subset of memory Ly49H^+^ NK cells with enhanced antiviral effector functions persists indefinitely (Sun et al., 2009). Similar types of adaptive NK cells develop in response to hapten sensitization (O’Leary et al., 2006), vaccinia virus infection (Gillard et al., 2011), and virus-like particle immunization of mice (Paust et al., 2010). Likewise, simian immunodeficiency virus-reactive memory NK cells develop in rhesus macaques after virus infection or immunization (Reeves et al., 2015). Collectively, animal studies point to existence of long-lived, virus-dependent subpopulations of memory NK cells that are likely better antiviral effectors than their naïve counterparts.

Several types of memory NK cells have been characterized in humans. These include memory NK cells induced by cytokines (Cooper et al., 2009), varicella zoster virus exposure (Nikzad et al., 2019), antibody-mediated stimulation (Zhang et al., 2013), or CMV-derived peptides (Hammer et al., 2018). High frequencies of NK cells expressing the activating receptor NKG2C are frequently observed in CMV seropositive individuals (Guma et al., 2006a). These NKG2C^+^ cells undergo proliferative expansion during primary CMV infection in transplant patients (Beziat et al., 2013) and in response to CMV-infected fibroblasts (Guma et al., 2004), IL-12-producing infected monocytes (Rolle et al., 2014), and CMV UL40-derived peptides (Hammer et al., 2018). CMV-associated adaptive NK cells expressing NKG2C display altered DNA methylation patterns and reduced expression of signaling molecules, including FcRγ, spleen tyrosine kinase (SYK), and EWS/FLI1-associated transcript 2 (EAT-2) (Lee et al., 2015; Schlums et al., 2015). These FcRγ^neg^, SYK^neg^, and/or EAT-2^neg^ NK cells also generally lack expression of the transcription factor promyelocytic leukaemia zinc finger protein (PLZF) (Schlums et al., 2015). These phenotypic changes are linked to more potent antibody-dependent activation, expansion, and function of these adaptive NK cells relative to other NK-cell subsets. NK cells with reduced expression of FcRγ, SYK, or EAT-2 are also detected in CMV seronegative individuals, with a minor fraction (10%) of individuals displaying significant expansions of this population (Schlums et al., 2015).

The crucial function of NK cells in immune defense against CMV coupled with the discovery that distinct subsets of NK cells emerge after infection, collectively suggest that targeted induction of these subsets of NK cells during immunization may provide enhanced protection against CMV infection. The capacity of existing vaccines to elicit transient or sustained expansion of CMV-associated human NK cells has not been reported. In this study, we interrogate longitudinal peripheral blood mononuclear cell (PBMC) samples collected from MF59-adjuvanted CMV glycoprotein B (gB) vaccine or placebo recipients who locally participated in clinical trial NCT00133497 (Bernstein et al., 2016). Our study reveals vaccine-independent oscillation of FcRγ^neg^ NK cell frequencies, but not those of NKG2C^+^ NK cells, in the blood of CMV seronegative individuals. Phenotypic and functional characterization of FcRγ^neg^ NK cells in this CMV seronegative cohort reveals distinct features from those reported for FcRγ^neg^ NK cell in individuals infected with CMV. These finding provoke re-evaluation of the paradigm concerning NK-cell subset dynamics in humans.

Several types of memory NK cells have been characterized in humans. These include memory NK cells induced by cytokines [33], varicella zoster virus exposure [34], antibody-mediated stimulation [35], or CMV-derived peptides [36]. High frequencies of NK cells expressing the activating receptor NKG2C are frequently observed in CMV seropositive individuals [37]. These NKG2C^high^ cells undergo proliferative expansion during primary CMV infection in transplant patients [38] and in response to CMV-infected fibroblasts [39], IL-12-producing infected monocytes [40], and CMV UL40-derived peptides [36]. CMV-associated adaptive NK cells expressing NKG2C display altered DNA methylation patterns and reduced expression of signaling molecules, including FcRγ, spleen tyrosine kinase (SYK), and EWS/FLI1-associated transcript 2 (EAT-2) [41, 42]. CMV-associated NK cell expansions with reduced expression of FcRγ, SYK, and/or EAT-2 generally lack expression of the transcription factor PLZF and may also express activating KIR or lack DAP12-coupled receptors altogether. NK cells with reduced expression of FcRγ, SYK, or EAT-2 are also detected in CMV seronegative individuals, with 10% of individuals displaying significant expansions of this population. These phenotypic changes are linked to more potent antibody-dependent activation, expansion, and function of these adaptive NK cells relative to other NK-cell subsets.

The crucial function of NK cells in immune defense against CMV coupled with the discovery that distinct subsets of NK cells emerge after infection, collectively suggest that targeted induction of these subsets of NK cells during immunization may provide enhanced protection against CMV infection. The capacity of existing vaccines to elicit transient or sustained expansion of CMV-associated human NK cells has not been reported. In this study, we interrogate longitudinal peripheral blood mononuclear cell (PBMC) samples collected from MF59-adjuvanted CMV glycoprotein B (gB) vaccine or placebo recipients who locally participated in clinical trial NCT00133497 [2]. Our study reveals vaccine-independent oscillation of FcRγ^neg^ NK cell frequencies, but not those of NKG2C^high^ NK cells, in human blood from CMV seronegative individuals. These finding provoke re-evaluation of the paradigm concerning NK-cell subset dynamics in humans.

## Materials and Methods

### CMV vaccine trial

This study was approved by the Cincinnati Children’s Hospital Medical Center Institutional Review Board and conducted by the Cincinnati Vaccine and Treatment Evaluation Unit (VTEU) as part of CMV vaccine trial NCT00133497. Study participants were 12- to 17-year-old healthy adolescent females confirmed CMV seronegative at the start of the study. Only samples from the Cincinnati site of this clinical trial were available for the purposes of the present study. Furthermore, only those subjects with available samples spanning trial duration were used for experimental analyses. As a result, a total of 40 participants were randomized into two groups (n=20/group) receiving either 3 doses of CMV gB subunit vaccine in MF59 adjuvant (20 μg gB and 10.75 mg MF59, Sanofi Pasteur) or sterile saline (Sodium chloride 0.9%) placebo by intramuscular injection in the deltoid on days 0, 30, and 180 of protocol (Bernstein et al., 2016). Urine, saliva and blood were collected throughout time course to assess CMV infection by PCR and seroconversion to non-vaccine CMV antigens, respectively. The 40 subjects evaluated longitudinally in the present study remained CMV negative throughout sampling period. Three additional vaccine trial participants who were part of the placebo group and became positive for CMV infection during longitudinal sampling period were used to examine NK-cell subset frequencies at time points subsequent to natural acquisition of CMV infection. Peripheral blood mononuclear cells (PBMC) were collected and cryopreserved at screening and various time points (days 0, 1, 30, 60, 180 and 210) of trial (Bernstein et al., 2016).

### NK-cell phenotypic analyses

PBMC were concomitantly stained and assessed by flow cytometry during a single experimental run (or block). A volunteer blood donor with a high percentage of NKG2C^+^ NK cells extraneous to vaccine trial was selected as a positive control for NKG2C staining and included in each block of vaccine trial participant samples to benchmark stain validity and reproducibility. Expression of FcRγ, SYK, and EAT-2 are benchmarked against CD4 T cells in the same sample, where the latter cells do not express these proteins (Schlums et al., 2015). Phenotypic analyses of PBMCs were performed using fluorochrome-conjugated antibodies. Cells were stained for surface markers using CD3 (OKT3, Biolegend), CD19 (HIB19, BD Biosciences), CD4 (RPA-T4, BD Biosciences), CD14 (M5E2, BD Biosciences), CD56 (N901, Beckman Coulter), NKG2C (REA205, Miltenyi Biotech), NKG2A (Z199, Beckman Coulter), CD57 (HCD57, Biolegend), CD16 (3G8, BD Biosciences), Ki-67 (Ki-67, Biolegend) and a fixable live-dead stain (Pacific Green, Invitrogen) in FACS buffer (HBSS supplemented with 5% fetal bovine serum and 2 µm EDTA). Following surface staining, cells were fixed in 2% paraformaldehyde (Fisher Scientific) and permeabilized with 0.04% Triton X-100 (Sigma Aldrich). Intracellular staining in FACS buffer with 2% bovine serum albumin was then performed to identify FcRγ (polyclonal rabbit, Millipore), EAT-2 (polyclonal rabbit, ProteinTech Group), SYK (4D10.1, eBioscience) markers. Intracellular EAT-2 staining was followed by secondary staining with polyclonal anti-rabbit IgG (Invitrogen).

### NK-cell functional analyses

PBMC samples were thawed rapidly in a 37°C water bath and cell number and viability were determined using 0.4% Trypan Blue (Thermo Fisher Scientific). Cells were cultured at 5×10^5^ per well in a 96 well U-shaped plate (Corning Life Sciences) at 37°C in 5% CO_2_. Control wells received only media (RPMI 1640 media (Thermo Fisher Scientific) supplemented with 10% fetal bovine serum), while cytokine-stimulated wells received a combination of IL-12 (10 ng/ml), IL-15 (100 ng/ml), and IL-18 (100 ng/ml) (Schlums et al., 2015). After 18 hours of culture, Golgi Plug (BD Biosciences) and Golgi Stop (BD Biosciences) were added for an additional 6 hours at final concentrations of 1 µg/ml and 2 µM, respectively. To assess antibody dependent cell cytotoxicity (ADCC), a third well of 5×10^5^ PBMC for each sample were mixed with 1.25×10^5^ P815 cells (2:1 effector to target (E:T) ratio) pre-incubated with 2.5 μg/ml anti-CD32 (Clone 2.4G2, Bio-X-Cell). Cells were incubated in the presence of Golgi Stop and Golgi Plug for a total of 6 hours (Hart et al., 2019). Anti-CD107a (H4A3, Biolegend) at 1:200 dilution was added to all cells in the final 6 hours of stimulation. Intracellular staining in FACS buffer was performed to assess IFN-γ (4S.B3, Biolegend) production. Flow cytometric data for all phenotypic and functional analyses were obtained using an LSR Fortessa instrument (BD Biosciences) and analyzed via FlowJo_v10 software (Treestar).

### T-distributed stochastic neighbor embedding (t-SNE) analyses

The tSNE algorithm of FlowJo_v10 was used to visualize dimensionality of NK cell subsets over time. For each donor, the data at individual time point was down sampled (gated on CD56^dim^ NK cells) and then concatenated to create three dimensionally reduced t-SNE plots. Populations expressing or lacking various proteins were overlaid on t-SNE plots to identify subset clusters.

### Statistical analyses

Differences between placebo and vaccine recipients were compared using mixed effects two way ANOVA with restricted maximum likelihood. Changes over time (0, 6, 7, 10, and 13 months) and treatment group (placebo and vaccine) in the proportion of CD56^bright^ and CD56^dim^ cells were evaluated using generalized linear mixed models with a Poisson distribution, log link function, and an offset of the logarithm of the total NK cell count specified. A random intercept and a random slope and an interaction term between time and treatment group was included in the model. Correlations between NK cell markers were determined by linear regression analysis. Differences in phenotype and function were determined by Student’s t-test and two-way ANOVA. Graphs were generated using GraphPad Prism and statistical tests were performed in Prism and SAS 9.4 (SAS Institute Inc., Cary NC).

## Results

### CMV gB vaccine trial cohort

To determine whether CMV vaccination strategies can trigger emergence of CMV-associated NK cell subsets, we examined a longitudinal series of PBMC from a subset (n=40, **Table 1**) of CMV vaccine trial participants (NCT00133497) for whom a full set of samples were available. Half of the study participants received three intramuscular injections of CMV gB in MF59 adjuvant while the placebo group was administered sterile saline in place of the vaccine (**Figure 1A**). Vaccine recipients exhibited a robust gB-specific antibody response (**Figure 1B**). None of the selected 40 study participants acquired CMV infection during the study period, as measured by PCR testing for CMV in urine and for seroconversion against non-vaccine CMV antigens (Bernstein et al., 2016).

**Table 1.** Patient Demographics. Age, race, and ethnicity of female subjects who received placebo or gB/MF59 vaccine (n=20 per group).

|  | gB/MF59 | Placebo | Total |
| --- | --- | --- | --- |
| # subjects | 20 | 20 | 40 |
| Age Category at Vaccination, n (%) |  |  |  |
| 12-15 years | 14 (70) | 11 (55) | 25 |
| 15-17 years | 6 (30) | 9 (45) | 15 |
| Race, n (%) |  |  |  |
| Black | 7 (35) | 7 (35) | 14 |
| Caucasian | 12 (60) | 11 (55) | 23 |
| Other | 1 (5) | 2 (10) | 3 |
| Ethnic Origin, n (%) |  |  |  |
| Hispanic/Latino | 1 (5) | 0 (0) | 1 |
| Not | 19 (95) | 20 (100) | 39 |

**Figure 1.**
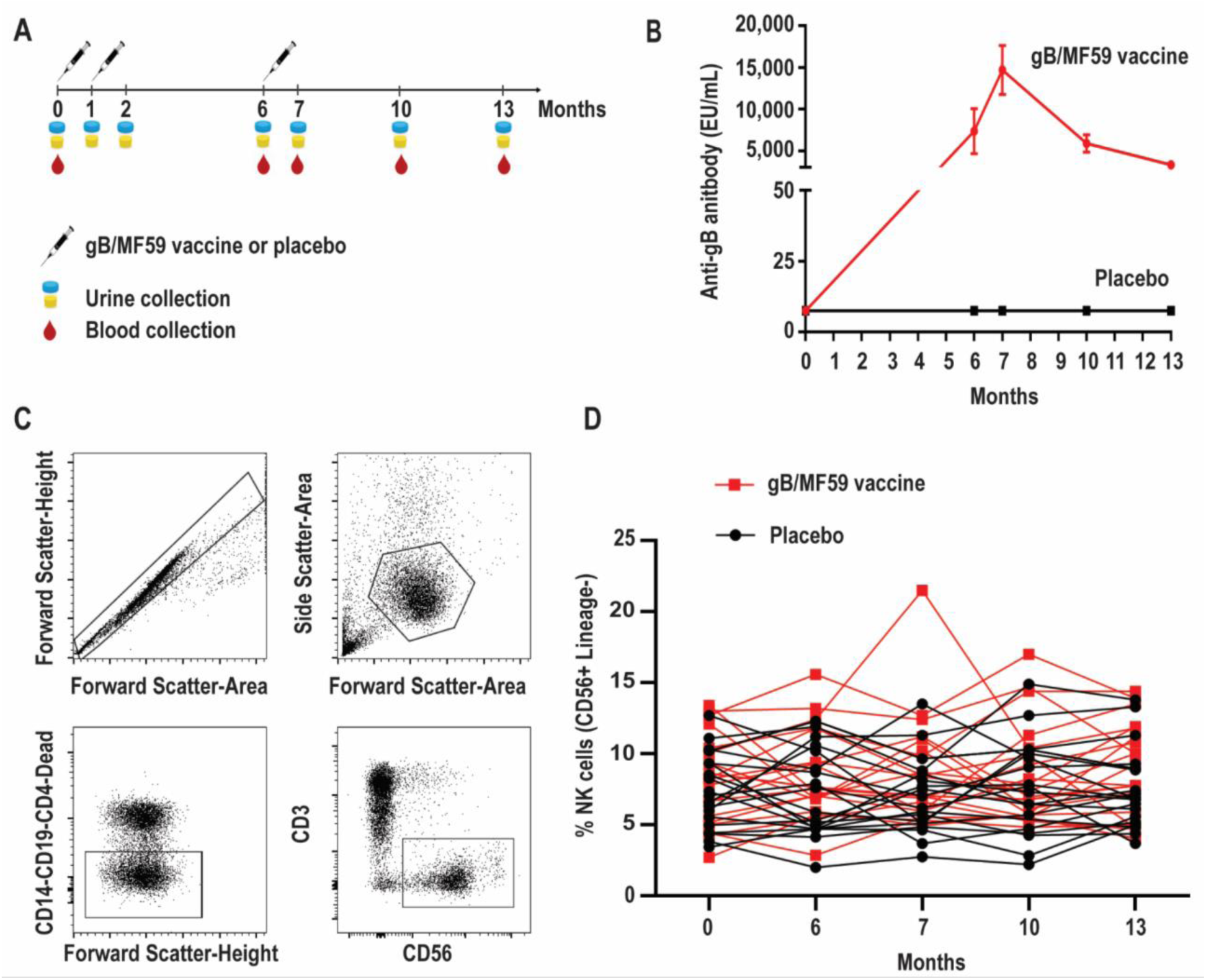
Longitudinal antibody and NK-cell responses of vaccine study participants. (A) Schematic representation of the timeline of vaccine trial depicting three administrations of gB/MF59 vaccine or placebo (sterile saline) and timing of urine and blood samples collection. (B) Sera anti-gB antibody titers for placebo (n=20) and vaccine (n=20) recipients analyzed in present study. (C) Representative flow cytometry gating of singlets, live lymphocytes, lineage-negative (CD19, CD14, CD4, CD3) CD56^+^ NK cells based on forward scatter (height and area), side scatter (area), viability dye uptake, and surface marker expression. (D) Resulting frequencies of gated NK cells in individual vaccine and placebo recipients over sampling period.

### Minimal variation in total NK-cell frequencies over time

We first assessed the proportion of total NK cells (CD56^+^ CD3^-^ CD19^-^ CD14^-^ CD4^-^) in PBMC. **Figure 1C** depicts the gating scheme used to identify NK cells in our samples. There was a broad range (2.0-21.5%) of NK-cell proportions across study participants (**Figure 1D**). The mean proportion of NK cells across all time points is similar in groups receiving placebo or vaccine (Placebo = 7.4%, Vaccine = 8.6%, p=0.16), while the changes in NK cell proportions over time between the placebo or vaccine group were not statistically significantly different (p=0.71).

NK cells can be stratified based on CD56 expression into CD56^dim^ and CD56^bright^ subsets (**Figure 2A**) that exhibit distinct phenotypic and functional characteristics (Ellis and Fisher, 1989). The CD56^dim^ subset comprises a mean 88.8±1.14% (average of all time points) of circulating NK cells in study participants (**Figure 2B**), where the ratio between CD56^bright^ and CD56^dim^ cells in vaccine and placebo groups is relatively consistent over study time points (**Figure 2C**). Specifically, time did not modify the effect between the placebo and vaccine groups regarding CD56^dim^ cell counts (p=0.38). In addition, neither time (p=0.97) nor treatment group (mean vaccine: 6.50% (95% CI: 5.34%, 7.91%); mean placebo: 5.51% (95% CI: 4.51%, 6.74%), p=0.24) were independently associated with CD56^dim^ cell counts. For CD56^bright^ cell counts, time modified the effect of placebo and vaccine groups (p=0.01); wherein CD56^bright^ count increased by a factor of 1.25 (95% CI: 1.07, 1.46, p=0.01) in the vaccine group but the change in the placebo group was not statistically significant (**Table 2**), which is consistent with other vaccine studies (Berthoud et al., 2009; Scott-Algara et al., 2010; Przemska-Kosicka et al., 2018).

**Table 2.** Change in NK cell CD56^bright^ and CD56^dim^ subsets over time. Mean (95% confidence interval) percentage of CD56^dim^ and CD56^bright^ cells by time and treatment group.

|  | Placebo Mean (95% CI) | Vaccine Mean (95% CI) |
| --- | --- | --- |
| CD56<sup>Dim</sup> (interaction p-value = 0.38) |  |  |
| 0 m | 5.95% (4.73%, 7.50%) | 6.20% (4.95%, 7.76%) |
| 6 m | 5.41% (4.29%, 6.81%) | 6.34% (5.07%, 7.94%) |
| 7 m | 5.15% (4.09%, 6.49%) | 6.99% (5.58%, 8.75%) |
| 10 m | 5.41% (4.30%, 6.82%) | 6.45% (5.15%, 8.07%) |
| 13 m | 5.67% (4.49%, 7.16%) | 6.53% (5.21%, 8.20%) |
| CD56<sup>Bright</sup> (interaction p-value = 0.01) |  |  |
| 0 m | 0.62% (0.50%, 0.77%) | 0.58% (0.47%, 0.72%) |
| 6 m | 0.61% (0.49%, 0.76%) | 0.60% (0.48%, 0.74%) |
| 7 m | 0.60% (0.49%, 0.75%) | 0.69% (0.56%, 0.86%) |
| 10 m | 0.55% (0.44%, 0.68%) | 0.70% (0.57%, 0.87%) |
| 13 m | 0.59% (0.47%, 0.73%) | 0.72% (0.58%, 0.90%) |

**Figure 2.**
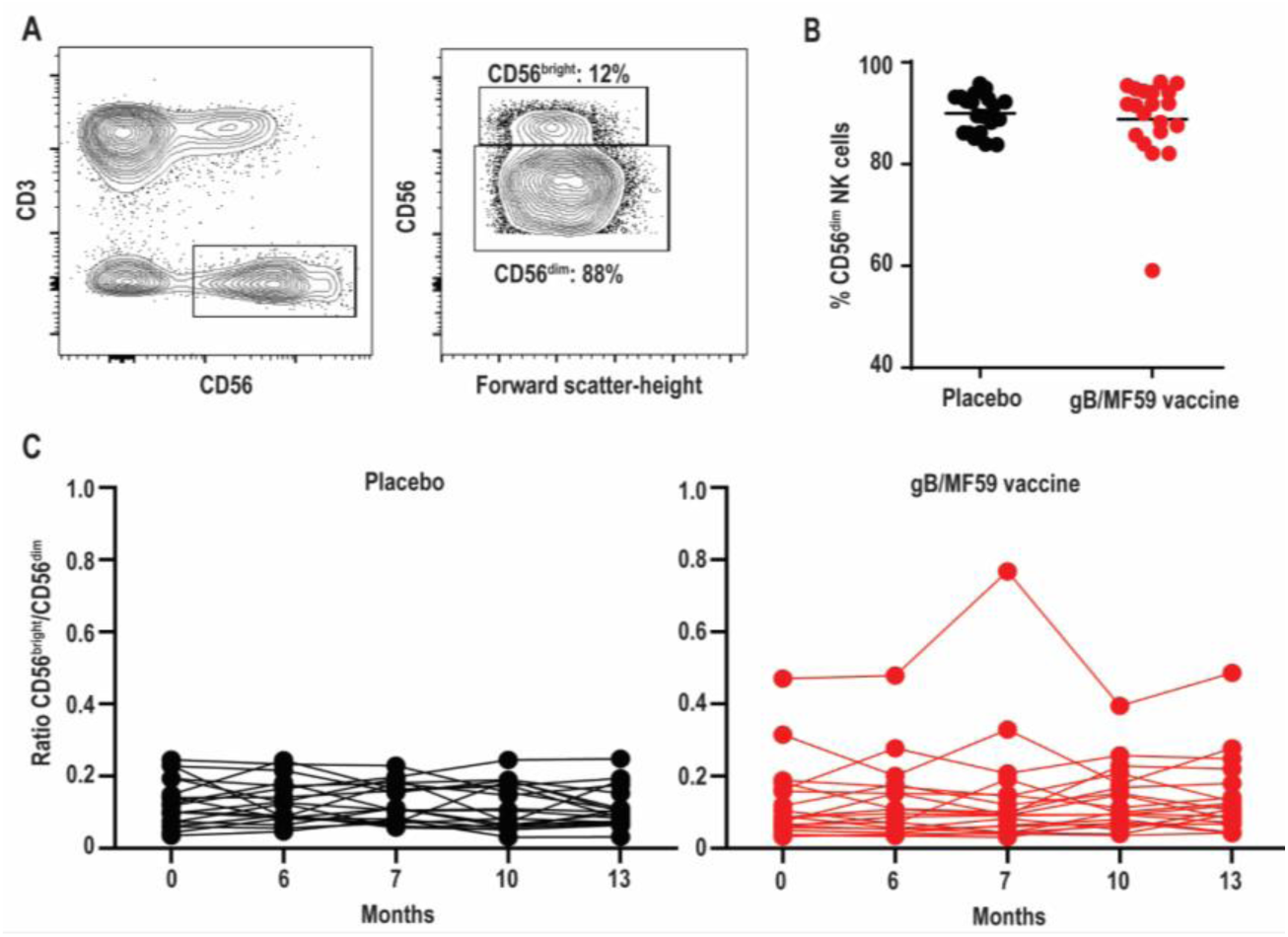
Stable proportions of CD56^dim^ and CD56^bright^ NK cells over time. (A) Representative gating of CD56^bright^ and CD56^dim^ events within the Lineage^-^ CD3^-^CD56^+^ NK cell gate, and (B) percentage of CD56^dim^ NK cells in each study participant averaged across all time points. Bar represents mean among treatment group (n=20/group). (C) Ratio of CD56^bright^ to CD56^dim^ NK cells in each study participant across vaccine trial time points. Statistically significant changes in subset proportion over time evaluated using generalized linear mixed model as described in Methods, with results presented in **Table 2**.

### Absence of CMV-dependent NKG2C^+^ *NK-cell subset*

High frequencies of NKG2C-expressing NK cells have been almost exclusively observed in CMV seropositive individuals (Guma et al., 2006a). This subset expands after CMV reactivation in organ or tissue transplant recipients (Beziat et al., 2013), and reflects activation of this subset by CMV UL40-derived peptides coupled with pro-inflammatory cytokines (Hammer et al., 2018). Due to the confirmed CMV negative status of vaccine trial participants throughout the duration of the vaccine study and the absence of UL40 antigens in the vaccine formulation, we hypothesized that NKG2C^+^ NK cell frequencies would be low at all time points. Using a positive control PBMC sample known to contain NKG2C^+^ NK cells (Schlums et al., 2015), we confirmed that our staining protocol can effectively detect this subset (**Figure 3A**). As expected, NKG2C^+^ NK cells were largely undetectable in all vaccine trial participants at baseline and the average absolute change in frequency from baseline proportions of this subset hardly varied across time in both placebo (0.046-0.41% range of mean absolute change from baseline visit) and vaccine (−0.83-0.96% range of mean absolute change from baseline visit) recipients (**Figure 3B**). Analysis of additional samples from vaccine trial participants (n=3) at time points after natural acquisition of CMV infection (confirmed by PCR & seroconversion to non-vaccine CMV antigens) remained negative for NKG2C^+^ NK cells over the study timeframe (data not shown).

**Figure 3.**
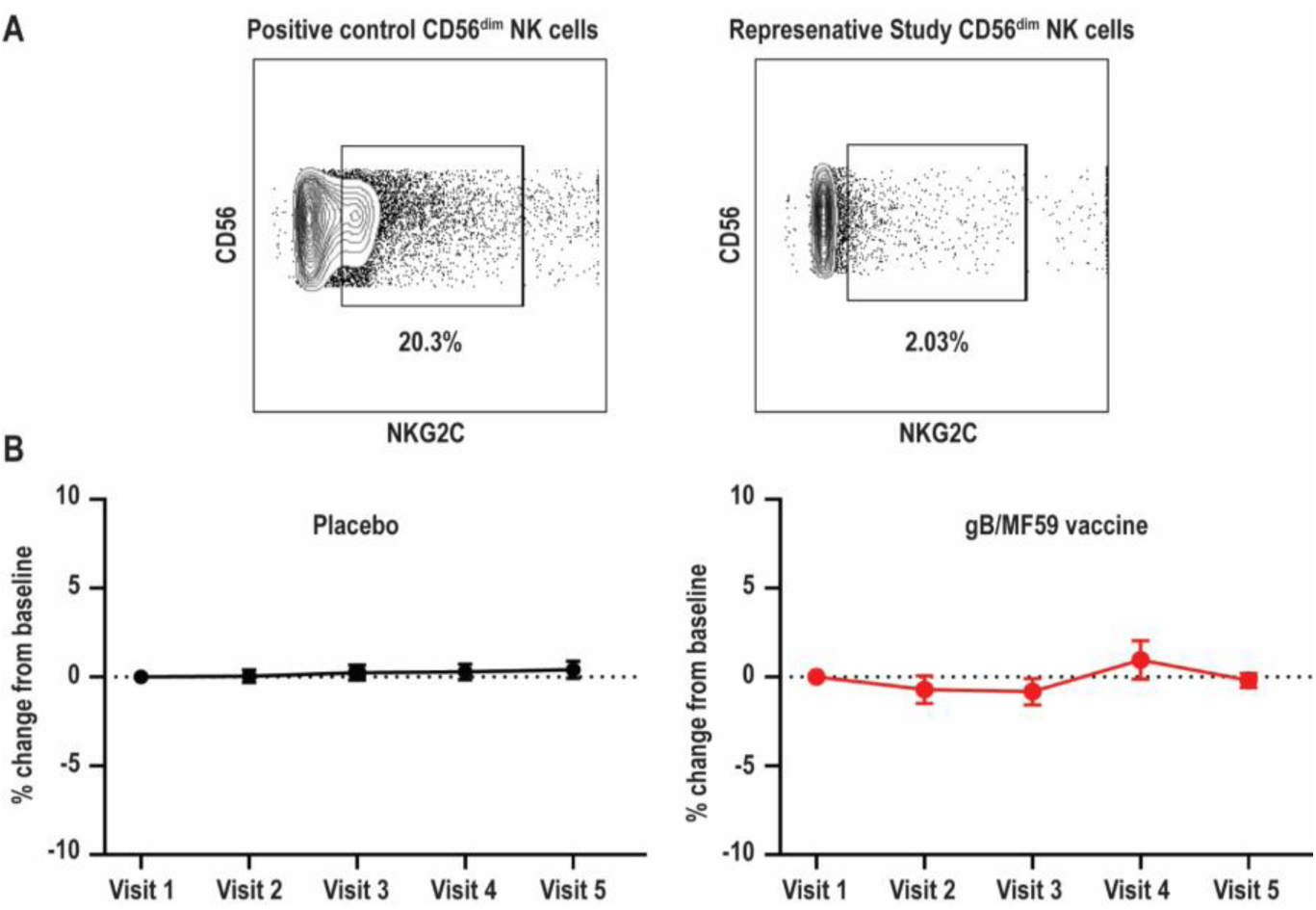
Absence of NKG2C^high^ NK cells in vaccine trial cohort. (A) Flow cytometry gating of NKG2C^high^ events among gated Lineage^-^CD56^dim^ NK cells in positive control sample and negligible staining for NKG2C on NK cells in a representative vaccine study participant. (B) For each study participant, the absolute change in proportion of NKG2C^high^ NK cells over time relative to measurement at baseline is calculated and presented as average (± standard error of the mean) of treatment group (n=20/group).

### CMV- and vaccine-independent dynamic changes in NK-cell subset frequencies

Expanded subsets of NK cells that lose expression of FcRγ, EAT-2, and/or SYK are expanded in approximately half of CMV seropositive individuals but can also be observed in seronegative donors, albeit less commonly (≤10% of individuals) and at much lower frequencies (Lee et al., 2015; Schlums et al., 2015). As these subsets can expand upon Fc receptor engagement by antibody (Zhang et al., 2013; Lee et al., 2015), we hypothesized that robust antibody responses against vaccine antigens may trigger accumulation of these subsets after vaccine prime and boost administration. We could detect NK cells within the CD56^dim^ subset that exhibited loss of FcRγ expression (**Figure 4A**). These FcRγ^neg^ NK cells concomitantly lacked EAT-2 and SYK expression in most study participants relative to their FcRγ^+^ NK cell counterparts (**Figure 4A**).

**Figure 4.**
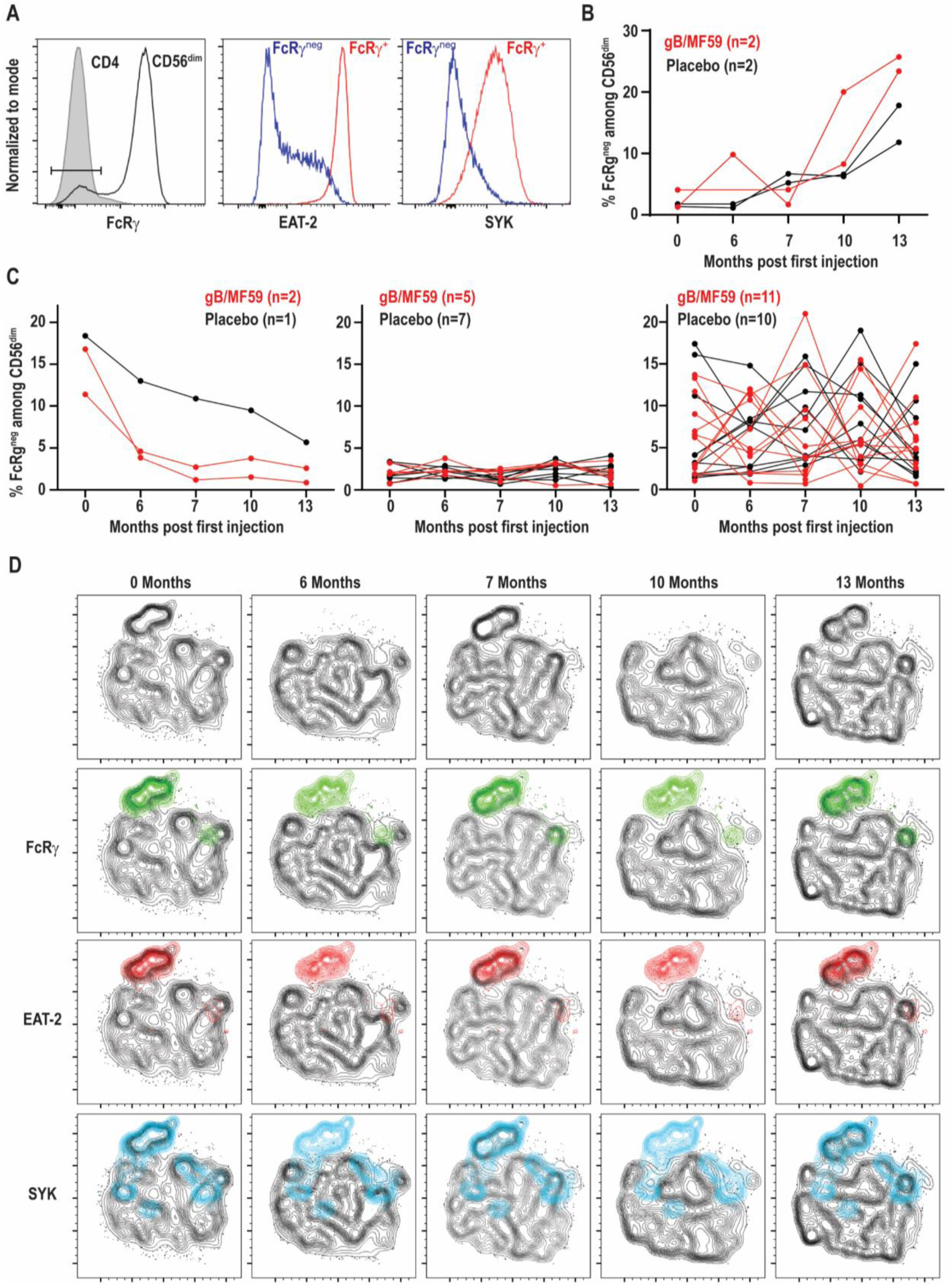
Dynamic vaccine-independent changes in NK-cell subset representation within repertoire over time. (A) Representative gating of FcRγ^neg^ CD56^dim^ NK cells (open histogram) relative to CD3^+^CD4^+^ T cells (shaded histogram) in the same sample. EAT-2 and SYK expression on gated FcRγ^neg^ (blue histogram) and FcRγ^+^ (red histogram) subsets of CD56^dim^ NK cells. (B) Proportions of FcRγ^neg^ CD56^dim^ NK cells over time in a subset of gB-MF59 (red) or placebo (black) recipients revealing expansion of these cells within the repertoire. (C) Proportions of FcRγ^neg^ CD56^dim^ NK cells over time in remaining study participants grouped based on pattern of subset contraction (left), absence of subset (middle), or ebb-and-flow changes in repertoire. (D) Location of each adaptive NK-cell subset in t-SNE distribution of a single study participant CD56^dim^ NK cell repertoire over time course is highlighted in green (FcRγ^neg^), red (EAT-2^neg^), and blue (SYK^neg^).

Interestingly, we detected a progressive increase in the frequency of FcRγ^neg^ NK cells following prime and boost immunization in a subset of vaccine recipients (**Figure 4B**). However, a similar pattern was observed in some placebo recipients. Moreover, the majority of individuals given vaccine (n=11) or placebo (n=10) exhibited transient elevations and depressions in the frequency of FcRγ^neg^ NK cells over time (**Figure 4C**). A smaller number of individuals in both groups demonstrated FcRγ^neg^ NK cells at baseline that disappeared over time, or lacked this subset of cells entirely (**Figure 4C**). High-dimensional analysis with t-SNE confirmed FcRγ^neg^, EAT-2^neg^, and SYK^neg^ NK cells largely cluster as one subset, the frequency of which changes over time within the selected study participant (**Figure 4D**).

### Variations in frequencies of FcRγ^neg^ NK cells are not associated with proliferation

In addition to increases and decreases in the proportion of FcRγ^neg^ populations among NK cells, the frequency of these cells among total blood leukocytes shows similar patterns of expansion and contraction (**Figure 5A**). The marked increases in the number of FcRγ^neg^ NK cells in some individuals over time are potentially attributable to periods of proliferative expansion. In fact, adaptive subsets of NK cells that accumulate during acute CMV infection of solid organ transplant recipients are characterized by heightened expression of Ki-67, an indication that these cells are highly proliferative (Lopez-Verges et al., 2011). Analysis of Ki-67 expression over time in a subset of six vaccine trial participants exhibiting ebb-and-flow representative of FcRγ^neg^ NK-cell within the NK-cell repertoire revealed relatively stable, low-level expression of Ki-67 on FcRγ^neg^ NK cells (**Figure 5B**). There was no clear visual relationship between Ki-67 expression and changes in frequency of FcRγ^neg^ NK cells among blood leukocytes (**Figure 5B**), and linear regression analysis of all time points analyzed revealed absence of significant relationship between the proportion of Ki-67-expressing FcRγ^neg^ NK cells and the fraction of NK cells that are FcRγ^neg^ (**Figure 5C**). Thus, within the limitations of these measurements and our sampling intervals, our data provide little evidence in support for the hypothesis that variations in FcRγ^neg^ NK cell frequencies are attributable to proliferative expansions of these cells.

**Figure 5.**
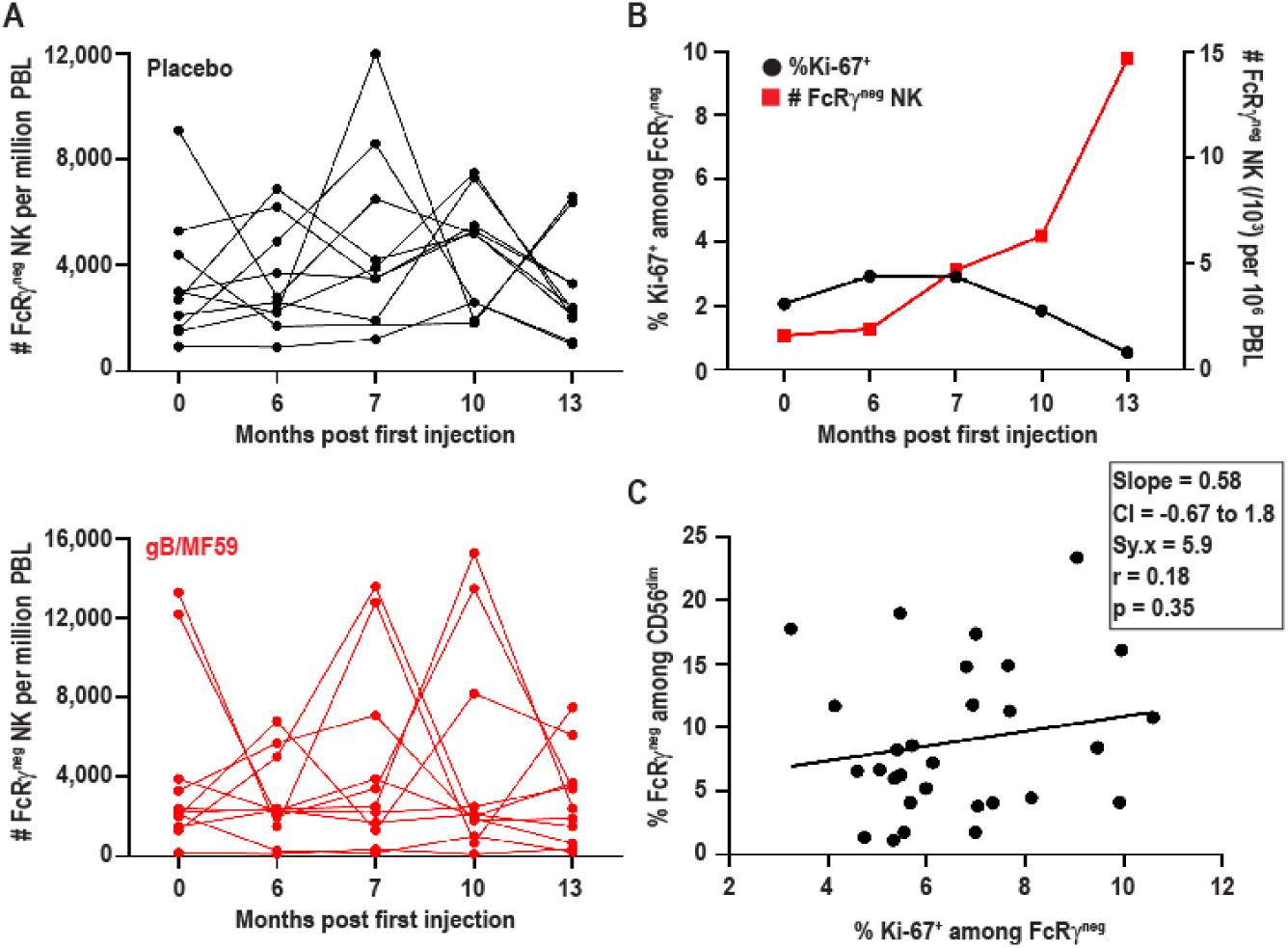
FcRγ^neg^ NK cell subset expansions in absence of increased Ki-67. (A) Frequencies of FcRγ^neg^ CD56^dim^ NK cells among PBL over time in a subset of gB-MF59 (red) or placebo (black) recipients exhibiting ebb-and-flow changes in repertoire. (B) Relationship between percent of FcRγ^neg^ NK cells staining Ki-67^+^ and expansion of FcRγ^neg^ NK cell subset over time. One representative individual is shown from among six vaccine trial participants with marked fluctuations in FcRγ^neg^ NK cell frequencies that were analyzed in this experiment. (C) Regression analysis of the linear relationship between the proportions of Ki-67^+^ cell within the FcRγ^neg^ NK cell subset and total FcRγ^neg^ NK cells. Slope, confidence interval (CI), residual standard error (Sy.x), correlation (r), and significant deviation of slope from zero (p) are shown.

### Distinct CD57 expression on FcRγ^neg^ NK cells in absence of CMV

While the proportions of FcRγ^neg^ EAT-2^neg^ SYK^neg^ NK cells vary among individuals and at different time points, the percentage of NK cells expressing other receptors associated with CMV infection, including CD57 (range 10-54%) or NKG2A (range 20-84%), exhibited little variation across time (**Figure 6A**). In fact, no statistically significant differences in CD57 (p=0.96) or NKG2A (p=0.75) expression were observed over time between placebo and vaccine groups. The temporally stable but heterogeneous expression of CD57 and NKG2A among individuals in the present study is consistent with prior observations (Ivarsson et al., 2017). As CMV infection is associated with increased expression of CD57 and down-regulation of NKG2A (Bjorkstrom et al., 2010), most notably among FcRγ^neg^ (Zhang et al., 2013) and NKG2C^high^ (Lopez-Verges et al., 2011) NK cells, the expression of these receptors was examined on the NK-cell subsets in vaccine trial participants (**Figure 6B**). FcRγ^neg^ NK cells detected in CMV-negative individuals in this study segregated as NKG2A^low^ relative to FcRγ^+^ cells (**Figure 6B**), consistent with previous studies (Zhang et al., 2013; Schlums et al., 2015). However, FcRγ^neg^ NK cells were not enriched for expression of the maturation marker CD57 (**Figure 6B**). In fact, the totality of FcRγ^neg^ NK cells observed across time points and individuals in this study expressed less CD57 than their FcRγ^+^counterparts (**Figure 6C**). Thus, FcRγ^neg^ NK cells are more prevalent in the NK-cell repertoire in this longitudinally examined study cohort and frequently exhibit dynamic changes in frequency over time as well as distinct CD57 expression patterns relative to FcRγ^neg^ NK cells in CMV infected individuals.

**Figure 6.**
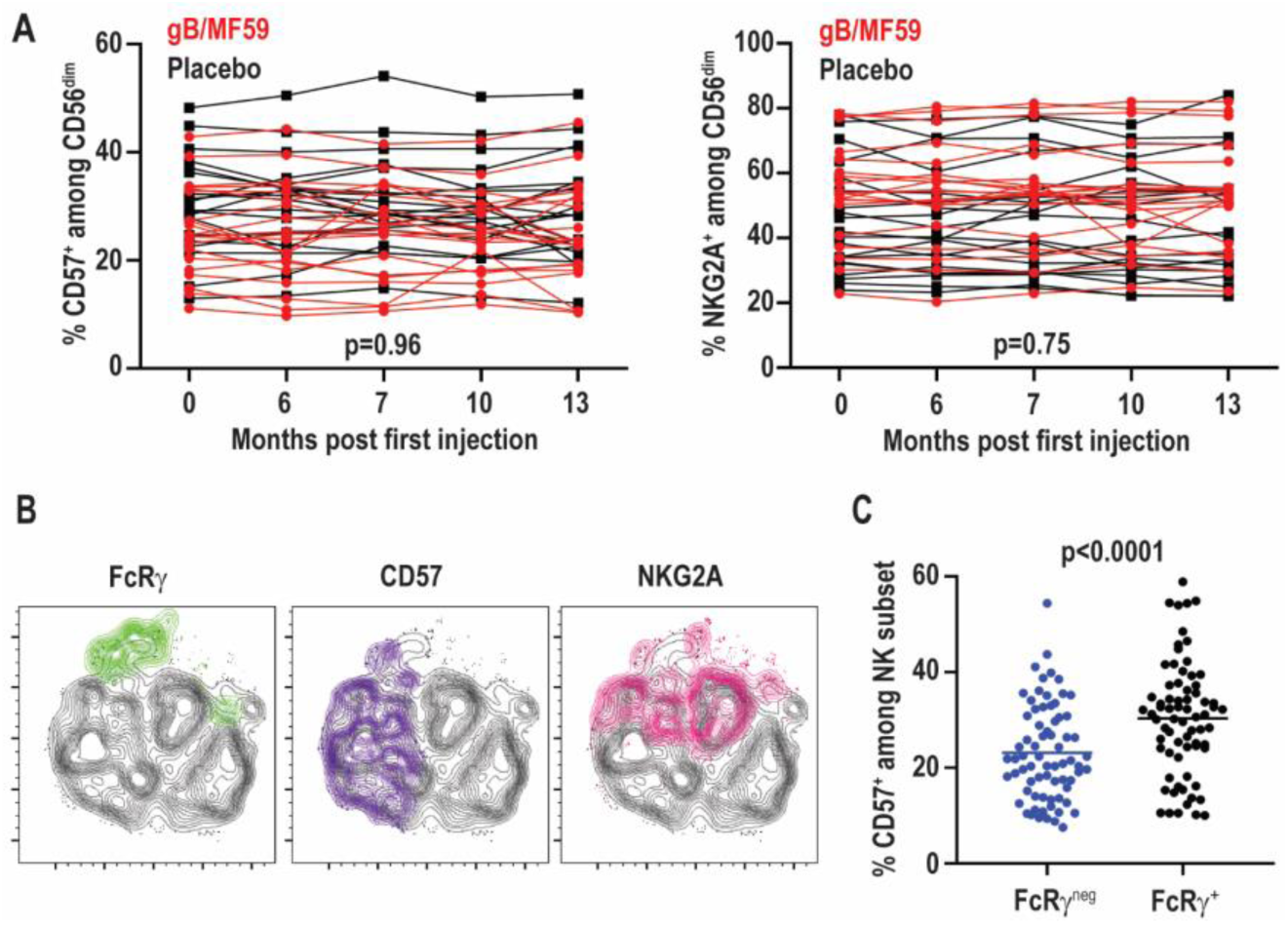
No change in CD57 and NKG2A over time. (A) Proportions of CD57^+^ and NKG2A^+^ CD3^-^CD56^dim^ NK cells in individual gB/MF59 vaccine (red) and placebo (black) recipients over time. Mixed effects two way ANOVA with restricted maximum likelihood was used to compare mean differences over time between both placebo and vaccine groups for each marker. (B) Representative location of FcRγ^neg^ (green), CD57^+^ (purple), and NKG2A^+^ (pink) cells at single time point in t-SNE distribution of a single study participant CD56^dim^ NK cell repertoire. (C) Expression of CD57 on FcRγ^neg^ and FcRγ^+^ CD56^dim^ NK cells across all time points and study participants where the FcRγ^neg^ subset was detectable (≥ 5% of CD56^dim^ NK cells). Statistically significance differences between repeated measures determined by Student’s t-test.

### No functional impact of FcRγ *loss in CMV seronegative subjects*

Signaling alterations of adaptive NK cells in CMV positive individuals lead to distinct functional capacities as compared to conventional NK cells (Hammer and Romagnani, 2017). In particular, FcRγ^neg^ NK cells in CMV infected individuals exhibit reduced IFN-γ production in response to cytokine stimulation (Schlums et al., 2015), but elevated antibody-dependent effector function (Zhang et al., 2013). Functional responses of FcRγ^neg^ NK cells in the absence of CMV were examined at a total of 18 samples from a subset of six vaccine trial participants scoring positive for FcRγ^neg^ NK cells. In contrast to observations in CMV seropositive individuals, IL-12 and IL-18 cytokine stimulation did not lead to statistically significant differences in IFN-γ production (p=0.41) or degranulation as measured by CD107a exposure (p=0.67) between FcRγ^neg^ and FcRγ^+^ NK cells in the CMV seronegative vaccine cohort (**Figure 7A**). FcRγ^neg^ and FcRγ^+^ NK cells also exhibited comparable degranulation (p=0.58) and IFN-γ production (p=0.38) when stimulated with P815 cells pre-incubated with α-CD16 antibody (**Figure 7A**). Of note, FcRγ^neg^ NK cells in vaccine recipients produced slightly more IFN-γ but displayed similar degranulation in response to α-CD16-bound P815 relative to the same cells in individuals receiving placebo (data not shown).

**Figure 7.**
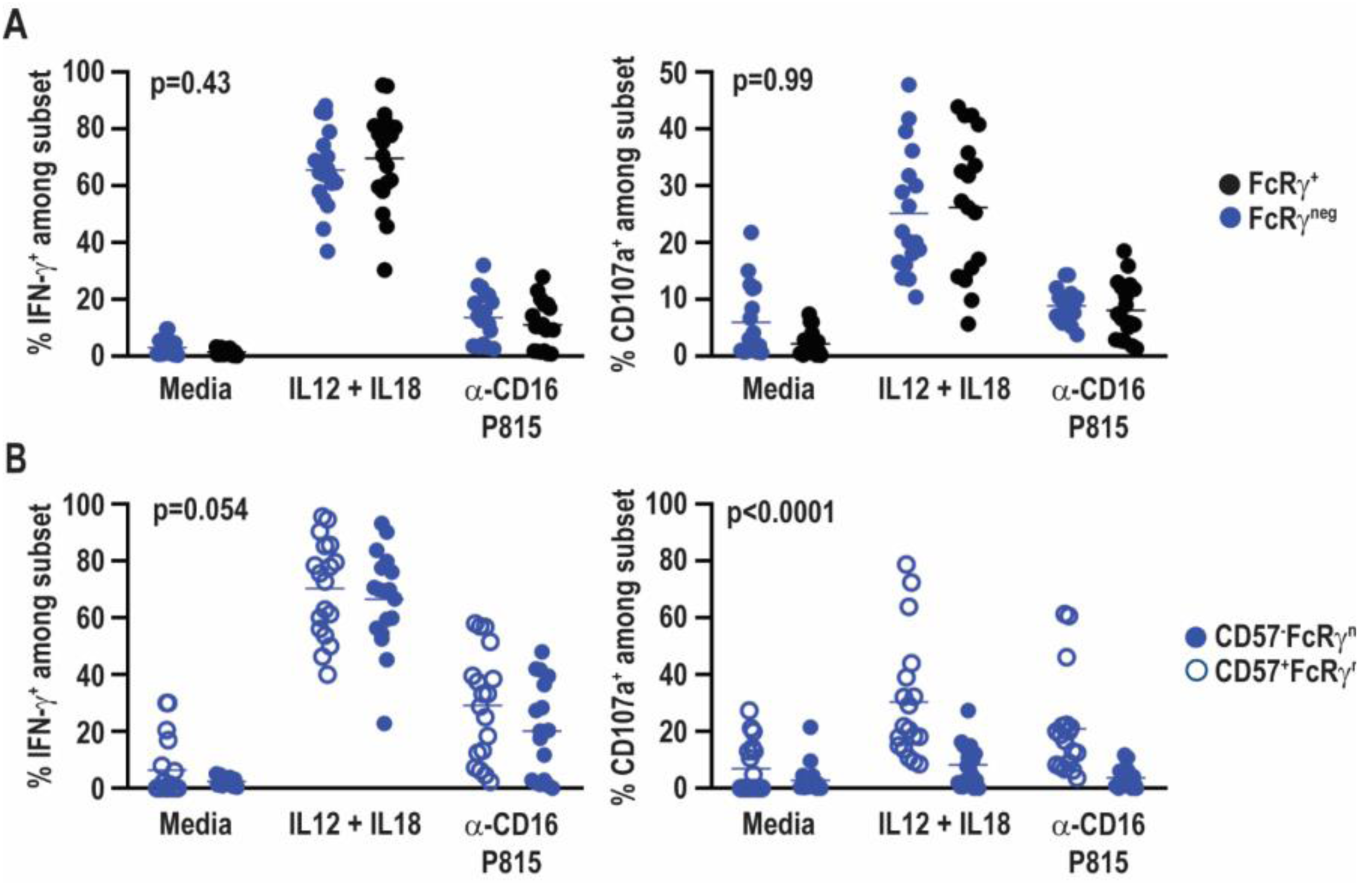
No functional impact of FcRγ deficiency in HCMV-negative individuals. Eighteen samples from six vaccine trial participants (3 gM/BF59 and 3 placebo) scoring positive for FcRγ^neg^ NK cells were stimulated with either IL-12+IL-18 for 24 hours or P815 cells labeled with α-CD16 antibody for 6 hours. Degranulation and IFN-γ production assessed by addition of fluorochrome-labeled α-CD107a antibody as well as GolgiPlug and GolgiStop during final 6 hours of incubation. Proportions of CD107a^+^ and IFN-γ^+^ events among (A) FcRγ^neg^ (blue) and FcRγ^+^ (black) CD3^neg^ CD56^dim^ NK cells or (B) CD57^+^ (open circles) and CD57^neg^ (closed circles) FcRγ^neg^ NK cells. Statistical significant differences between groups was determined by two-way ANOVA.

CD57^+^ NK cells are also differentially sensitive to cytokine and antibody-dependent stimulation compared to CD57^neg^ NK cells (Lopez-Verges et al., 2010). FcRγ^neg^ NK cells in the present vaccine cohort exhibit a distinct CD57 expression pattern compared to FcRγ^neg^ NK cells in CMV seropositive individuals (Bjorkstrom et al., 2010; Zhang et al., 2013). Therefore, functional responses of CD57^neg^ FcRγ^neg^ and CD57^+^ FcRγ^neg^ NK cells were compared within CMV seronegative vaccine trial participants. CD57^neg^ and CD57^+^ FcRγ^neg^ NK cells exhibited similar IFN-γ production after stimulation with IL-12 and IL-18 (p=0.51) or α-CD16 antibody bound P815 cells (p=0.14) (**Figure 7B**). However, CD57^+^ FcRγ^neg^ NK cells degranulated more robustly than their CD57^neg^ FcRγ^neg^ NK cell counterparts in response to either cytokine or P815+α-CD16 stimulation (**Figure 7B**).

## Discussion

Past cross-sectional analyses suggest that adaptive subsets of NK cells are rarely present in the absence of CMV infection, whereas the frequencies of these adaptive NK cells are markedly increased in the majority of CMV seropositive individuals (Lopez-Verges et al., 2011; Della Chiesa et al., 2012; Muntasell et al., 2013; Lee et al., 2015; Schlums et al., 2015). The present longitudinal analysis of NK cells in healthy CMV-negative individuals affirms the lack of NKG2C-expressing NK cells in CMV-naïve persons (Schlums et al., 2015), yet challenges the paradigm that the FcRγ^neg^ NK cell subset is infrequent or absent in CMV seronegative individuals. In contrast to the current prototype, the majority (28 of 40, 70%) of the healthy, demonstrably CMV-negative adolescent women profiled in this clinical study exhibit measurable frequencies of FcRγ^neg^ NK cells during at least one study time point, with dynamic changes in the frequency of these cells among circulating NK cells over time. Changes in frequencies of these subsets did not correlate with vaccine administration or vaccine-antigen-specific antibody titers (data not shown), suggesting that undefined environmental factors promote oscillations in the representation of these subsets among total circulating NK cells.

The low frequencies of NKG2C-expressing NK cells across all time points in the forty vaccine trial participants is consistent with absence of CMV infection of these individuals and the purported link between CMV gene products and expansion of this subset of NK cells (Guma et al., 2006b; Wu et al., 2013; Luetke-Eversloh et al., 2014). Likewise, we observed very low (0.22% ± 0.15% of live lymphocytes) but highly stable frequencies of infection-associated CD56^neg^ CD16^+^ NK cells in our cohort (data not shown), consistent with absence of CMV and other viruses linked to this unusual NK-cell population. Within the limitations of our sampling scheme, our results support the hypothesis that CMV gB and MF59 are insufficient to stimulate differentiation or accumulation of NKG2C^+^ NK cells. Since CMV UL40-derived peptides presented by HLA-E are critically required for HCMV-driven NKG2C expansion (Hammer et al., 2018), incorporation of UL40 into next generation vaccines may more effectively elicit NKG2C^+^ memory NK cell expansion.

In contrast to both the tight link between CMV and NKG2C^+^ NK cells and the reported rarity of FcRγ^neg^ NK cells in CMV-seronegative individuals, the present longitudinal data suggest that the latter NK cell subset may be commonly present in some NK-cell repertoires and can exhibit dynamic changes in frequency. There was no correlation between numeric increases in FcRγ^neg^ NK cells and expression of the proliferation marker Ki-67, suggesting that release of this subset from tissues may be a greater factor in these dynamic changes than proliferative expansion. However, the timing of experimental sampling in the present study likely precludes precise determination of a link between proliferation and FcRγ^neg^ NK cell accumulation. The majority (28 of 40, 70%) CMV seronegative individual in our study exhibited populations of FcRγ^neg^ NK cells greater than 10% during at least one of the five time points analyzed over a year-long study period. These data contrast a previous cross-sectional studies which found expansions of NK cells lacking FcRγ, EAT-2, and/or SYK in 6 out of 69 CMV seronegative adults (Schlums et al., 2015). The fraction of study participants scoring positive for FcRγ^neg^ NK cell subsets at any given time point in our study ranged from 30% to 45%, suggesting that additional factors may distinguish the two study populations. Moreover, the FcRγ^neg^ NK cells measured in this study appear to differ from those observed in CMV-infected individuals with regards to expression of the maturation marker CD57 (Lopez-Verges et al., 2010; Lopez-Verges et al., 2011; Zhang et al., 2013; Schlums et al., 2015).

In addition to their distinct phenotype, the FcRγ^neg^ NK cells measured here differ in their functional activity as compared to their counterparts in CMV positive subjects (Schlums et al., 2015). Namely, the FcRγ^neg^ NK cells in the present study exhibit similar capacity to make IFN-γ and degranulate as FcRγ^+^ NK cells in responses to cytokines or antibody-dependent stimuli. We speculate that CMV-independent FcRγ^neg^ NK cells are unlikely to bear hypomethylation at the *IFNG* locus as a consequence of CMV infection (Schlums et al., 2015). As CD57 expression on NK cells is putatively linked to increased cytolytic potential, decreased sensitivity to inflammatory cytokines, and reduced proliferative potential (Nielsen et al., 2013), this phenotypic disparity of FcRγ^neg^ populations of NK cells in the absence of CMV may reflect important functional distinctions as well. The observed increase in degranulation of CD57^+^ FcRγ^neg^ relative to CD57^+^FcRγ^neg^ NK cells we observe consistent with the notion that CD57^+^ cells are more differentiated and have a distinct transcriptional signature in comparison to CD57^-^ NK cells (Lopez-Verges et al., 2010).

A major distinction of the present study population is the restriction to analysis of adolescent females. The influence of puberty-associated hormones and other pediatric variables on adaptive NK cell subsets is unknown. Therefore, it is possible that the present longitudinal study reveals dynamics of NK cell subsets that are unique to adolescents, or even adolescent females, that are not shared by adult CMV seronegative populations. Of note, NK cells express the alpha and beta estrogen receptors (ERα and ERβ) and exhibit function alterations in response to estrogen (Nilsson and Carlsten, 1994; Hao et al., 2008). Moreover, while KIR, CD57, and NKG2A expression on NK cells remains stable across menstruation cycles (Ivarsson et al., 2017; Feyaerts et al., 2018), the stability of the FcRγ^neg^ NK cell subset in this setting is less clear. Therefore, increased prevalence of CMV-associated NK cells or dynamic variation in the frequencies of the cells may reflect hormonal changes or environmental influences that are unique to or more common in adolescent females.

Besides these differences in age and gender of our study population, the participants in the CMV vaccine trial also exhibited a greater degree of racial diversity than was represented in previous cross-sectional studies (Schlums et al., 2015). Specifically, 35% of our vaccine trial participants were Black (i.e. African American). Although race assuredly impacts the NK-cell repertoire in the context of highly polymorphic receptors, including killer-cell immunoglobulin-like receptors (KIR), the effects of race on CMV-reactive NK cells and receptors associated with these subsets are less well defined. Intriguingly, 100% (15 of 15) of Black study participants demonstrated detectable FcRγ^neg^ NK cells at one or more time points of study, whereas only 58% (14 of 24) of Caucasian study participants exhibited FcRγ^neg^ NK cells in their repertoire. Thus, gender, race, genetics, and local environmental factors may all contribute to the distinct observations of adaptive NK cell frequencies in our study.

A key unanswered question concerns the nature of the stimuli provoking longitudinal changes in frequency of NK cell subsets. A recent study of barcoded hematopoietic cells in rhesus macaques noted significant fluctuations in the clonal composition of NK cells over time (Wu et al., 2018). Our study stringently controlled for CMV exposure via urine and blood analyses (Bernstein et al., 2016). Moreover, the results do not support a relationship between CMV gB vaccination or gB-specific antibody titers and altered frequencies of NK cell subsets. The present results contrast with marked change in NK-cell phenotype and function observed following protein subunit or inactivated virus vaccine administration in CMV seropositive individuals (Long et al., 2008; Scott-Algara et al., 2010; Nielsen et al., 2015; Goodier et al., 2016; Darboe et al., 2017). Nonetheless, other subclinical acute infections, vaccinations (e.g. seasonal influenza vaccine), inflammatory events, environmental exposures (i.e. allergens), or shifts in microbiota composition could alter the composition of the NK-cell repertoire. We speculate that these environmental stimuli or associated immune responses (i.e. antibody elaboration) provoke the expansion, differentiation, or release of FcRγ^neg^ NK cells into the circulation. The elevated frequency of these NK cell subsets in CMV-positive individuals may reflect an altered tempo or magnitude of these natural oscillations, or a greater regularity of the instigating stimulus. Alternatively, as the frequencies of these subsets appear to be more stable in CMV-seropositive individuals (Beziat et al., 2013), aspects of the inflammatory environment during chronic CMV infection may more efficiently maintain these populations. Given that the MF59 adjuvant used in this CMV vaccine is designed for optimal stimulation of T and B-cell responses, future studies aimed at ascertaining the nature of inflammatory cues promoting adaptive NK cells will yield key insights into the types of adjuvants that may be applied to intentional promote sustained expansion of these NK cell subsets in next generation vaccines.

Our results, to our knowledge, represent the first longitudinal study of CMV-associated NK-cell subsets in healthy CMV seronegative individuals. Here, we had the unique ability to gain insight into the intra-individual variation in the frequency of NK-cell subsets following gB/MF59 vaccination. We show that the lack of change in NKG2C expression was consistent with absence of CMV infection, confirming the stringent association of this virus with NKG2C^+^ NK cells. However, we also present evidence suggesting that presence of FcRγ^neg^, EAT-2^neg^, and SYK^neg^ NK cells in the repertoire may be more temporally dynamic and CMV-independent than previously thought. These data also reveal potentially important functional differences between CMV independent FcRγ^neg^ NK cells and those accumulating in the context of CMV infection. Future work examining age and gender related differences as well as longitudinal analyses of post-transplant patients may give further insight into the variegated expression of CMV-associated NK-cell subsets.

## Acknowledgements

This research was supported by the Cincinnati Children’s Research Foundation and the National Center for Advancing Translational Sciences of the National Institutes of Health (NIH) under Award Number UL1 TR001425. Investigators on this project are supported by NIH grants AI125413 (L.A.), DA038017, AI148080 and AR073228 (S.N.W.). The original clinical trial was supported by NIH contract HHSN272200800006C (D.I.B.). The Cincinnati Children’s Flow Cytometry Core is supported by NIH grants AR047363, AR070549, DK078392, DK090971, S10OD025045 and S10OD023410. D.O. is supported by a fellowship from the American Heart Association. Y.T.B. is supported by European Research Council under the European Union’s Seventh Framework Program (FP/2007-2013)/ERC Grant Agreement no. 311335, the Swedish Research Council, Norwegian Research Council, Swedish Foundation for Strategic Research, Wallenberg Foundation, Swedish Cancer Foundation, Swedish Childhood Cancer Foundation, and the Stockholm County Council and Karolinska Institutet Center for Innovative Medicine. We would like to thank G. Hart (U. Minnesota) for insights into the staining protocol for FcRγ^neg^ EAT-2^neg^ SYK^neg^ NK cells

